# DnaA and SspA Regulation of the *iraD* gene of *E. coli:* an alternative DNA damage response independent of LexA/RecA

**DOI:** 10.1101/2021.05.20.444958

**Authors:** Thalia H. Sass, Alexander E. Ferrazzoli, Susan T. Lovett

## Abstract

The transcription factor RpoS (σS) of *Escherichia coli* controls a large number of genes important for tolerance of a variety of stress conditions. IraD promotes the post-translation stability of RpoS by inhibition of RssB, an adaptor protein for ClpXP degradation. We have previously documented DNA damage induction of *iraD* expression, independent of the SOS response. Both *iraD* and *rpoS* are required for tolerance to DNA damaging treatments such as H_2_O_2_ and the replication inhibitor azidothymidine in the log phase of growth. Using luciferase gene fusions to the 672 bp *iraD* upstream region, we show here that both promoters of *iraD* are induced by AZT. Genetic analysis suggests that both promoters are repressed by DnaA-ATP, partially dependent on a putative DnaA box at −81 bp, and regulated by RIDA (<u>r</u>egulatory <u>i</u>nactivation of <u>D</u>na<u>A</u>), dependent on the DnaN processivity clamp. By electrophoretic mobility shift assays we show that purified DnaA protein binds to the *iraD* upstream region, so DnaA regulation of IraD is likely to be direct. DNA damage induction of *iraD* during log phase growth is abolished in the *dnaA*-T174P mutant, suggesting that DNA damage, in some way, relieves DnaA repression, possibly through the accumulation of replication clamps and enhanced RIDA. We demonstrate that the RNA-polymerase associated factor, SspA (<u>s</u>tringent <u>s</u>tarvation <u>p</u>rotein <u>A</u>), induced by the accumulation of ppGpp, also affects IraD expression, with a positive effect on constitutive expression and a negative effect on AZT-induced expression, in a fashion independent of DnaA.

**SIGNIFICANCE:** DNA damage can lead to cell death or genomic instability. Cells have evolved transcriptional responses that sense DNA damage and up-regulate tolerance and repair factors; the LexA/RecA-regulated SOS response in *E. coli* was the first example of such a system. This work describes an alternative DNA damage response, controlled by DnaA and the IraD post-translational regulator of RpoS. The cellular signals for this response, we propose, are empty replication processivity (β) clamps that accumulate at replication blocks. IraD expression is also regulated by stringent starvation protein, SspA, induced by nutrient deprivation. This SOS-independent DNA damage response integrates a signal of incomplete replication with starvation to modulate expression of genes that promote the completion of replication.

## INTRODUCTION

The IraD protein of *Escherichia coli* is a member of a group of “anti-adaptor” proteins [1] that control the post-translational levels of RpoS, an alternative sigma factor central to the general stress response [2]. RpoS induces transcription of approximately 10% of the genes in *E. coli* and promotes survival to a number of stresses, including those induced by starvation, near UV-irradiation, desiccation, acidic pH, reactive oxygen species, temperature and osmotic shock. RpoS is also essential for virulence of a number of pathogenic proteobacteria [3].

Transcription of *iraD* is induced following DNA damage [4] and its expression rapidly stabilizes RpoS by trapping the RssB adaptor protein for ClpXP proteolysis in an inactive conformation [1, 5]. After DNA damage, IraD induction leads to RpoS accumulation in the cell and induction of the RpoS regulon, even during early exponential growth phase [4]. Stabilization of RpoS by IraD is required for optimal survival to certain DNA damaging agents, including H_2_O_2_, phleomycin and azidothymidine [4]. RpoS is also naturally induced during the transition from exponential growth to stationary phase, in the absence of DNA damage, and IraD appears to be the major determinant of increased RpoS levels during this transition [4]. Growth phase regulation of IraD is mediated by positive regulation of *iraD* promoters by ppGpp, the signaling molecule that accumulates during the “stringent response” to nutrient deprivation. IraD transcription regulation therefore integrates two cellular signals: DNA damage and starvation. IraD is also negatively regulated at the translational level by CsrA-repression of an upstream leader peptide that is translationally coupled to IraD [6]. CsrA, an RNA binding protein, represses translation of a number of genes induced during the stringent response [7]. Since RpoS itself induces CsrA transcription [8], this provides a potentially rapid negative feedback loop that may aid recovery after transient stress conditions.

The DNA damage induction of the *iraD* gene is of particular interest because it occurs independent of the canonical DNA damage response pathway [4], the “SOS response”, controlled by the LexA and RecA proteins of *E. coli*. A number of other genes show similar SOS-independent induction by DNA damage, although the molecular nature of the inducing signals and regulators of this response are not currently understood (reviewed in [9]). The best documented members of this SOS-independent group are the *dnaA-dnaN-recF* operon [10], which encodes the initiator for replication (*dnaA*), the replication processivity beta clamp (*dnaN*) and the recombination mediator protein (*recF*), and the *nrdAB* operon [11], encoding the major aerobic nucleotide reductase required to sustain deoxynucleotide pools. Both operons encode replication or repair factors and are regulated transcriptionally by DnaA protein [12-14], which functions as a transcription factor in addition to its role as a replication initiator protein [15].

DnaA is a member of the AAA+ ATPase family and its binding to DNA has been best characterized at *E. coli oriC*, the origin of replication, where it regulates replication initiation. DnaA binds to multiple 9-bp high-affinity sites, known as DnaA boxes [16], to which it binds in both ATP- and ADP-bound states. ATP binding promotes the formation of an oligomeric helical structure of DnaA [17], which aids cooperative binding and spreading to additional nearby low affinity sites [18], which are not bound in the ADP state. This DnaA-bound state promotes the unwinding of DNA and the loading of the DnaB fork helicase [19]. The architectural DNA binding proteins, IHF and FIS, influence DnaA binding to *oriC*, positively and negatively, respectively [16].

In addition to regulating replication initiation, DnaA can act either as a transcriptional repressor or activator and can influence transcription termination [20]. DnaA binding to regulate transcription is presumed to reflect the same kinds of DNA interactions that it exhibits at *oriC* to control replication initiation. Although the most stringent DnaA boxes are defined as the sequence 5’ TTWTNCACA [21], DnaA can bind to much more degenerate sequences in vitro and in vivo and is influenced strongly by nearby sites (reviewed in [20]). For this reason, DnaA binding sites are impossible to predict based on sequence information alone and must be determined empirically [20]. In addition, although simple repression by DnaA is common, some genes show complex effects by DnaA. For example, the *nrdAB* promoter region is both positively and negatively regulated by DnaA [22], depending on DnaA’s concentration and nucleotide state.

Although overall levels of DnaA protein do not appear to be involved in regulation of replication initiation [23], levels of ATP-bound DnaA appear to correlate with initiation capacity [24]. A host of cellular mechanisms in *E. coli* appear to control the levels of free DnaA-ATP, both positively and negatively (reviewed in [25]). Acidic phospholipids [26] and interactions with specific sequences known as “DARS” (DnaA reactivating sites)[27] promote the exchange of ADP with ATP and the reactivation of DnaA for replication initiation. Free DnaA-ATP can be reduced through binding to *datA*, the <u>D</u>na<u>A</u> titration locus, which is rich in DnaA-ATP binding sites (reviewed in [25]). An additional negative regulatory mechanism is “RIDA” (<u>r</u>egulatory *i*nactivation of <u>D</u>na<u>A</u>) in which DNA polymerase-free replication clamps bind a homolog of DnaA, Hda, which interacts with DnaA to promote ATP hydrolysis (reviewed in [25]). These mechanisms, by influencing levels of DnaA-ATP, should also have an impact on the genes whose transcription is regulated by DnaA, although this has not been extensively investigated. RIDA appears to activate *nrdAB* transcription in vivo, consistent with a repressive effect of DnaA-ATP on its promoter [14].

In this work, we further investigate the regulation of the *iraD*. Mutations in factors that affect DnaA-ATP levels, such as certain alleles of DnaA and the DnaN processivity clamp, have effects consistent with DnaA-ATP acting as a repressor for *iraD* transcription. We demonstrate by electrophoretic mobility shift assays that DnaA binds to the *iraD* promoter region with a strength and cooperativity similar to that at the *nrdA* and *dnaA* promoters.

In addition, because *iraD* expression is induced by the stringent response through the signaling molecule, ppGpp [28], we also investigated the effects of SspA, “stringent starvation protein A”. In *E. coli*, SspA is induced by the stringent response and entry into stationary phase, although its expression is not affected by RpoS [29]. Originally identified as a RNAP-associated protein [30], SspA can act as a transcriptional activator [31]. SspA is of interest because it is widely conserved and regulates virulence genes in a number of gram-negative bacteria [31-37], often as a heterodimeric complex with an additional transcriptional co-activator. SspA is required for lytic growth of bacteriophage P1 in *E. coli* where, with phage-encoded co-activator Lpa, it promotes late transcription from promoter P*s* in vitro and in vivo [31]. In *E. coli*, SspA also up-regulates a cohort of acid-resistance genes in stationary phase [38]. Homodimeric SspA may regulate initiation and promoter clearance through interactions with σ70 [39, 40].

## RESULTS

The *iraD* upstream intergenic region is unusually large for *E. coli*, at 672 bp. Our previous work identified two transcriptional start sites for *iraD:* on at −417 (referred to as P1) and one at −137 (P2). P1 appears to be induced specifically during the transition to stationary phase, whereas P2 transcripts are apparent in all growth phases. Both appear to respond to ppGpp levels, showing reduced expression in *relA spoT* mutants and increased levels in mutants carrying “stringent” alleles of *rpoB* [28], which cause the transcriptional repertoire to resemble stringent cells even in the absence of ppGpp production.

To study *iraD* expression more easily we constructed a fusion of the entire *iraD* 672 bp upstream intergenic region to the *Photorhabdus luxCDEAB* operon, allowing us to monitor expression by light production during continuous growth of a culture. In addition, to examine the contributions of the *iraD* constitutive promoter, P2, while retaining similar surrounding sequence contexts, a mutation was made that substitutes a 6 bp EagI restriction site at the −10 sequence of P1, which eliminates the strong induction of *iraD* expression in late log phase. Likewise, expression from P1 only can be monitored with a similar 6 bp ApaI substitution at the −10 sequence of P2. Using a plate reader for luminescence, we monitor cultures for *iraD* expression during the course of growth from early exponential to stationary phase (Figure 1), with and without sublethal doses of the replication inhibitor azidothymidine (AZT) at time 0. As we had seen from Northern blots [4], expression in early log phase is dominated by P2 whereas P1 is responsible for the elevation in luciferase expression that occurs during the late log phase of growth. Expression from both promoters is increased by AZT, reaching a plateau at about 120 minutes after treatment. As we had shown previously, induction by AZT was evident even in *lexA3* strains, which are defective in mounting the SOS response. If anything, *lexA3* enhances expression after AZT for both P1 and P2. Constitutive expression from P1was also elevated in late log phase by *lexA3*. We think that this elevation can be explained by the longer persistence of damage in the absence of the SOS response.

**Figure 1.**
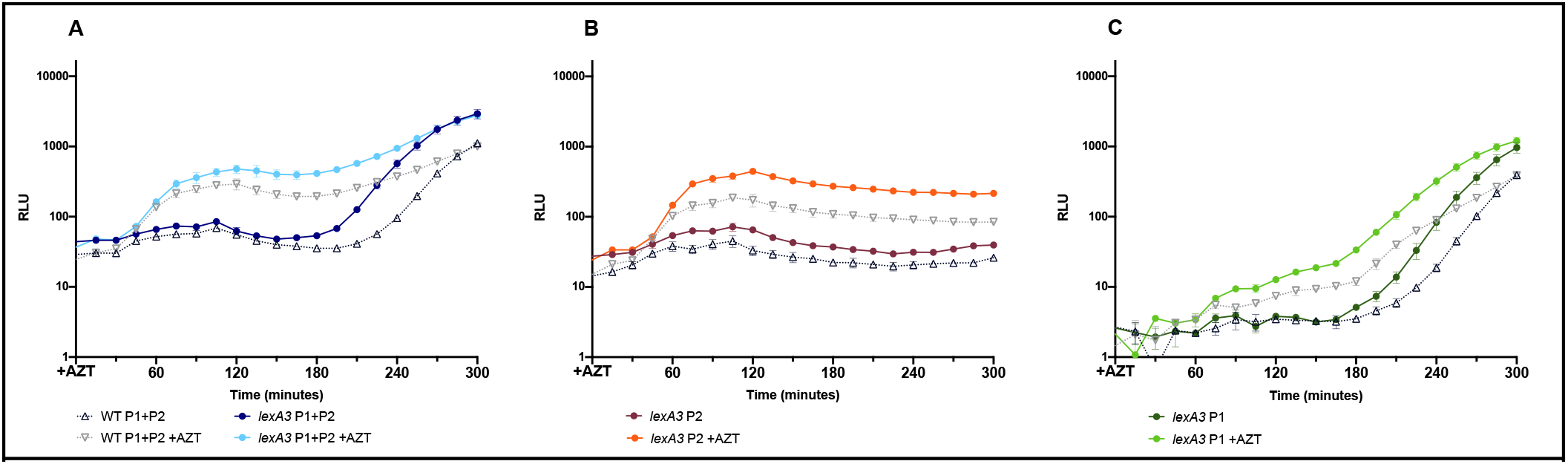
Luciferase expression from *iraD* promoter fusions, either from the wt upstream region (A), from P2 alone (B) or from P1 alone (C), with and without treatment with 12.5 ng/ml AZT at time 0. The wild-type expression is indicated with black open symbols and that in the *lexA3* genetic background with closed color symbols.

To determine if DnaA regulates *iraD* expression, we examined expression in strains that alter the relative amounts of DnaA-ATP vs. DnaA-ADP. We first compared luciferase expression in wild-type strains to a mutant in DnaA, *dnaA*-T174P, isolated [14] as a mutant that up-regulates *nrdAB* expression. In vitro, this mutation elevates the intrinsic ATPase of DnaA, thereby reducing the levels of DnaA-ATP in promoters, although more so from P2 (Figure 2A). We saw only a slight additional increase in expression after AZT treatment. We also examined a second, similar allele of *dnaA, dnaA* A345S, defective for ATP binding [14], and found that it similarly elevated constitutive expression of *iraD* at both promoters (Figure 2B). These results are consistent with DnaA-ATP acting as a repressor for *iraD* transcription at both P1 and P2.

**Figure 2.**
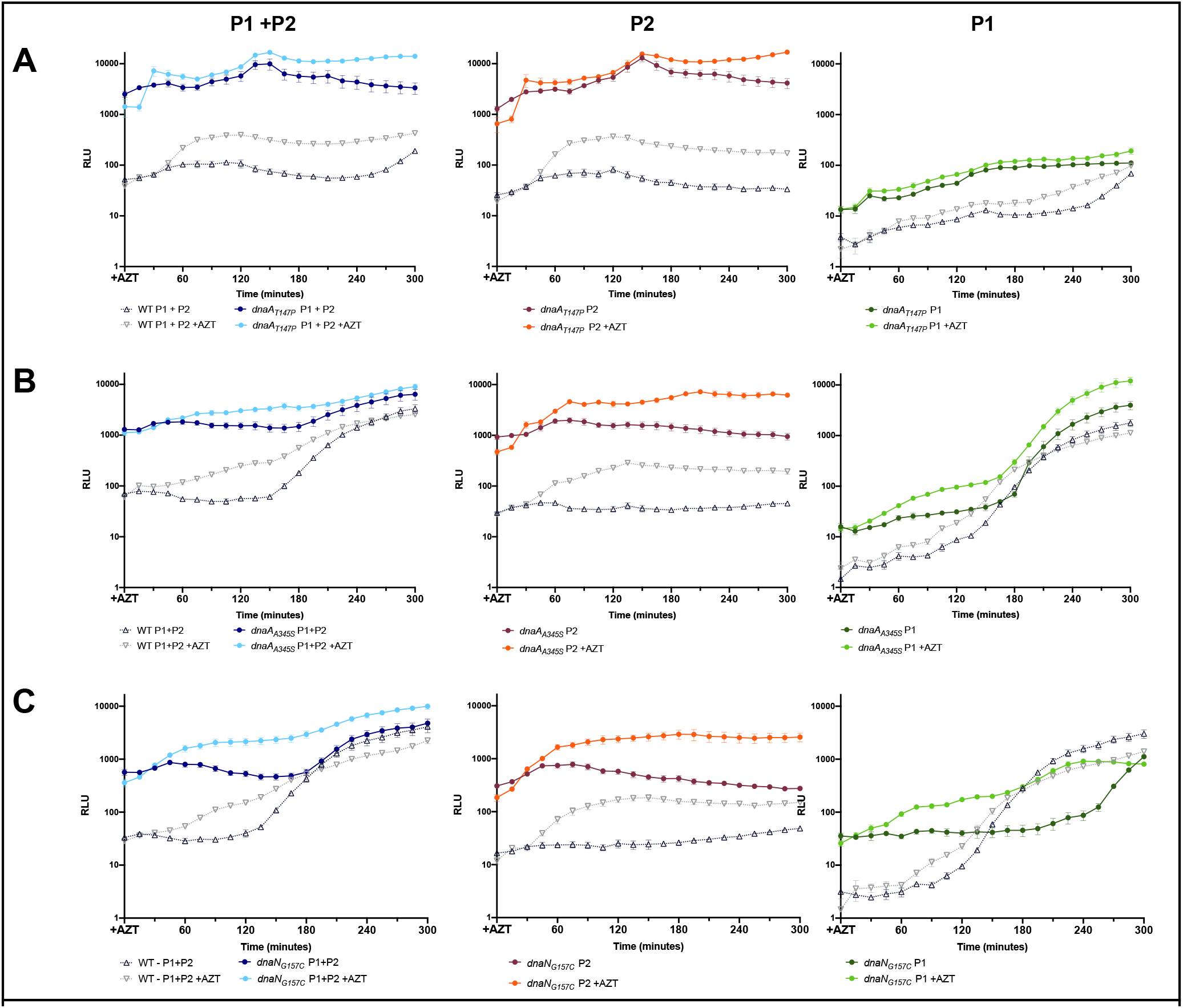
Luciferase expression from *iraD* promoter fusions in strains affecting RIDA. Data are shown either from the wt upstream region (left), from P2 alone (middle) or from P1 alone (right), with and without treatment with 12.5 ng/ml AZT at time 0. The wild-type expression is indicated with open symbols and that in the RIDA-genetic backgrounds with closed color symbols. A. dnaA-T174P strains, B. *dnaA*-A345S strains C. *dnaN*-G157C strains.

To determine if DnaA control of IraD had consequences for RpoS accumulation, we performed Western
blot analysis of RpoS in mid-exponential phase, in wild-type, *iraD*Δ and *dnaA*-T174P strains (Figure 3). As we had previously shown, levels of RpoS were reduced in *iraD* mutants. Strains carrying *dnaA*-T174Pshowed elevated RpoS levels, and this high level was partially *iraD*-dependent. This is consistent with the hypothesis that *dnaA*-T174P leads to elevated IraD and, subsequently, high RpoS levels. The fact that RpoS accumulation was not entirely IraD-dependent may indicate that DnaA is regulating other facets of RpoS control, potentially other Ira proteins [41], such as IraP or IraM.

**Figure 3.**
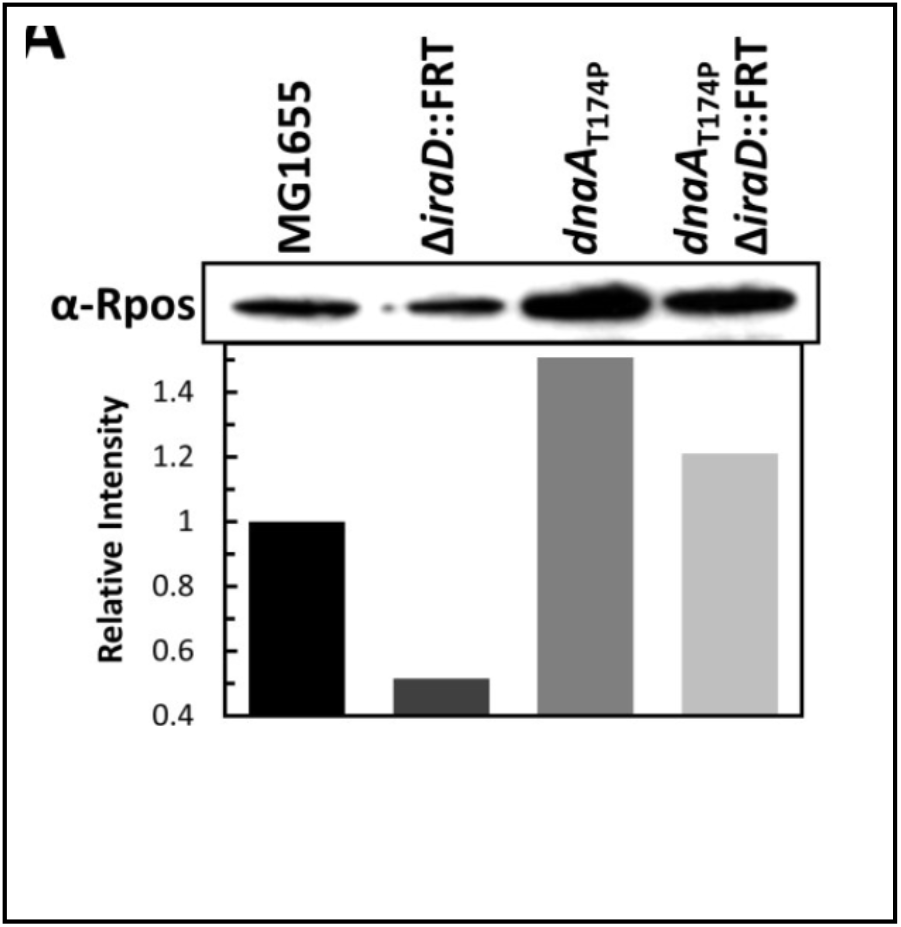
RpoS levels determined in Western blot in the indicated genetic backgrounds.

To determine if regulatory inactivation of DnaA, “RIDA”, affected regulation of *iraD*, we examined *iraD* expression using luciferase fusions in a mutant of the beta processivity clamp, DnaN. A <u>h</u>omolog of <u>D</u>na<u>A</u>, Hda protein, promotes hydrolysis of ATP by DnaA through its interaction with the replication processivity clamp, β, encoded by the *dnaN* gene. This hydrolysis, by depressing DnaA-ATP levels, down-regulates replication initiation at *oriC*. The *dnaN*-G157C allele, isolated as a mutant that increases *nrdAB* expression, exhibits an under-replication phenotype due to accumulation of DnaA-ADP. This is consistent with the *dnaN*-G157C allele promoting more regulatory inactivation of DnaA (RIDA), favoring DnaA-ADP over DnaA-ATP and relieving negative transcriptional regulation by DnaA-ATP. We observed that expression of *iraD*, as detected by luciferase expression, was elevated by *dnaN*-G157C, strongly in exponential phase and to a lesser extent upon entry to stationary phase (Figure 2C). Elevation of constitutive expression by *dnaN*-G157C was seen at P2 and in early log at P1, consistent with the notion that expression of *iraD* responds to the RIDA pathway. A depression of constitutive expression from P1 was seen in late log phase and the basis of this effect is not clear.

Although our data imply that DnaA-ATP negatively regulates *iraD*, these genetic effects could be direct, through DnaA binding to the *iraD* promoter region or, indirect, through DnaA control of other regulatory proteins. To test whether DnaA-ATP could bind to *iraD*, we performed electrophoretic mobility shift assays (EMSA) with purified DnaA protein at various concentrations and a DNA fragment encompassing the 672 bp *iraD* upstream region. DnaA did indeed bind the fragment, with an apparent Kd of 42 nM (Figure 4). Binding was observed to be cooperative, with a Hill coefficient (h) of 1.7.

**Figure 4.**
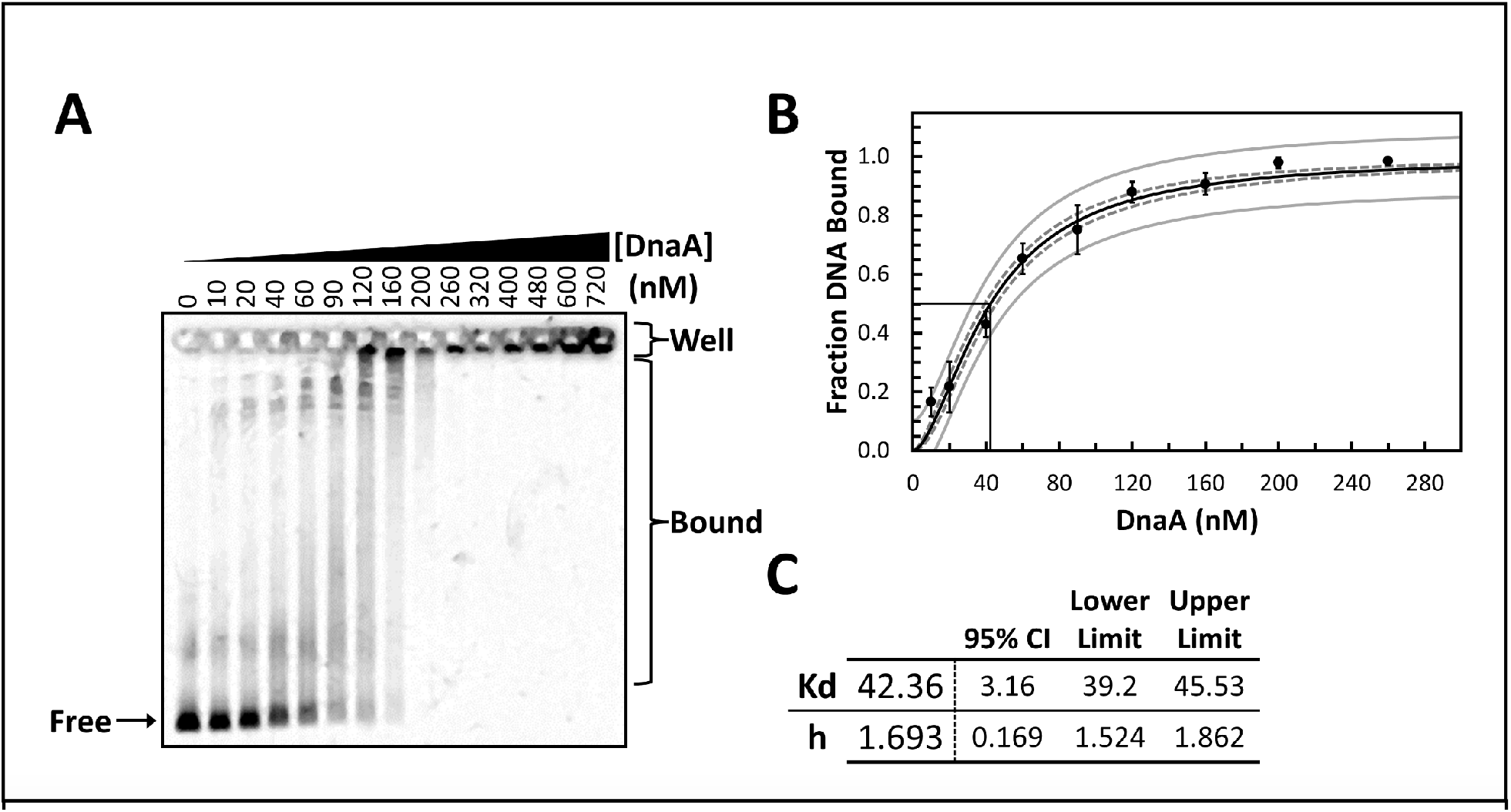
EMSA analysis of purified His-DnaA binding to the 672 bp *iraD* promoter region. A. Image of SYBR-Green DNA stained EMSA gel. Varying concentrations (0 – 720nM) of His-DnaA incubated with 10 nM PCR-generated DNA substrate for 10 minutes at 22°C. Under these electrophoretic conditions, DnaA protein does not migrate into the gel (Messer and Weigel 2003).B Binding isotherm generated from image densitometry of EMSA gels. Points are the average of four independent experiments. Error bars denote standard deviation. Black curve indicates the least-squares regression model from the compiled data from the four experimental experiments with the 95% confidence curve (dotted gray) and 95% prediction curve (solid gray) indicated. C. Summary table of parameters (Kd and Hill coefficient, h) obtained from the regression model with the 95% confidence interval and limits listed.

Analysis of the DNA shows that the strongest candidate for a potential DnaA box is at −81, relative to the ATG start codon of the open reading frame. This site (TTTTCCATA) matches 8/9 nucleotides for the stringent consensus for DnaA boxes (5’ TTWTNCACA) [21] and is a perfect match to the more relaxed consensus (5’ YYHTMCRHM, where Y=C/T; H= A/C/T; R=A/G and M=A/C)[42]. We mutated this putative DnaA box to CTCATTTTCCAT and measured expression from the intact upstream region and from P2. For both constructs Box −81 mutants showed consistently higher levels of expression (Figure 5), both constitutive and that induced by AZT, although to a lesser extent than was seen for the *dnaA* and *dnaN* alleles. The effects of Box −81 mutations are largely epistatic to the effects of DnaA-T174P (Figure 6), consistent with both alleles affecting repression of *iraD* expression by DnaA-ATP.

**Figure 5.**
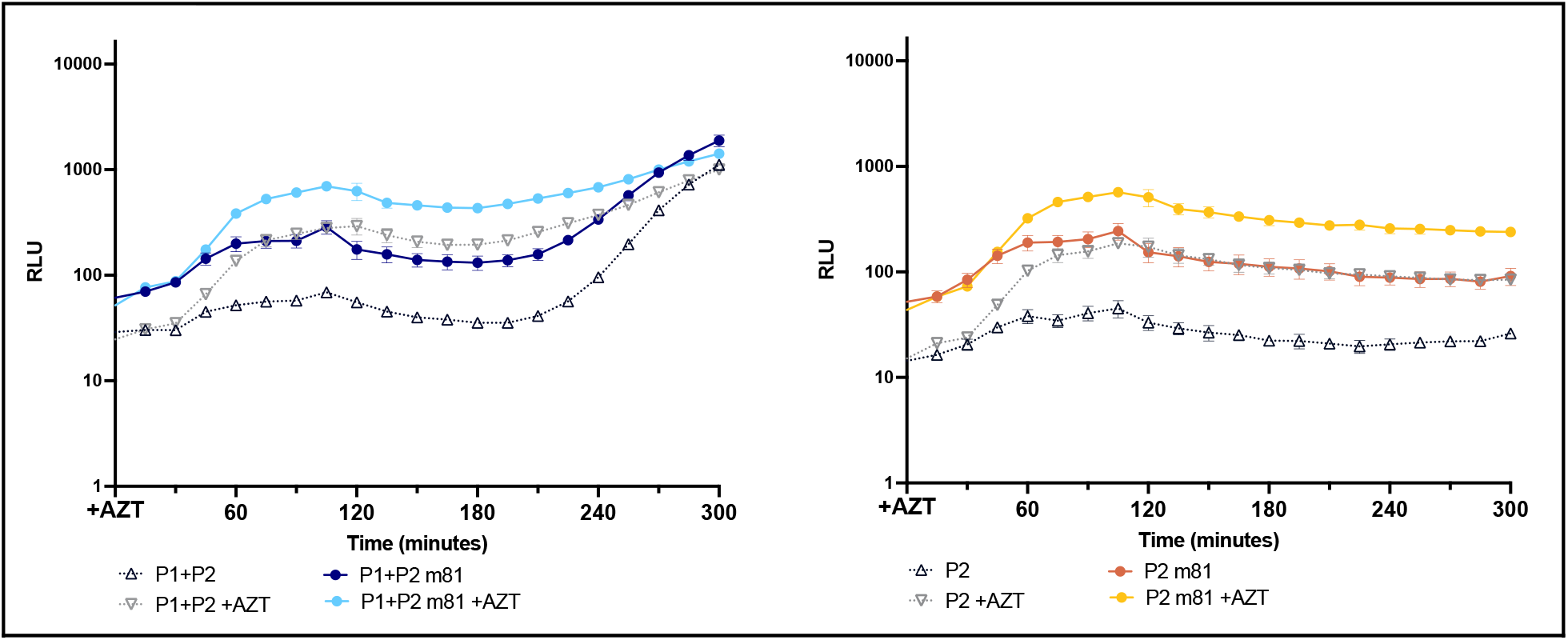
Effects of DnaA box −81 mutations on iraD expression from both promoters (left) or P2 only (right). Black indicates the wt iraD promoter expression and in color, expression from the box −81 mutants.

**Figure 6.**
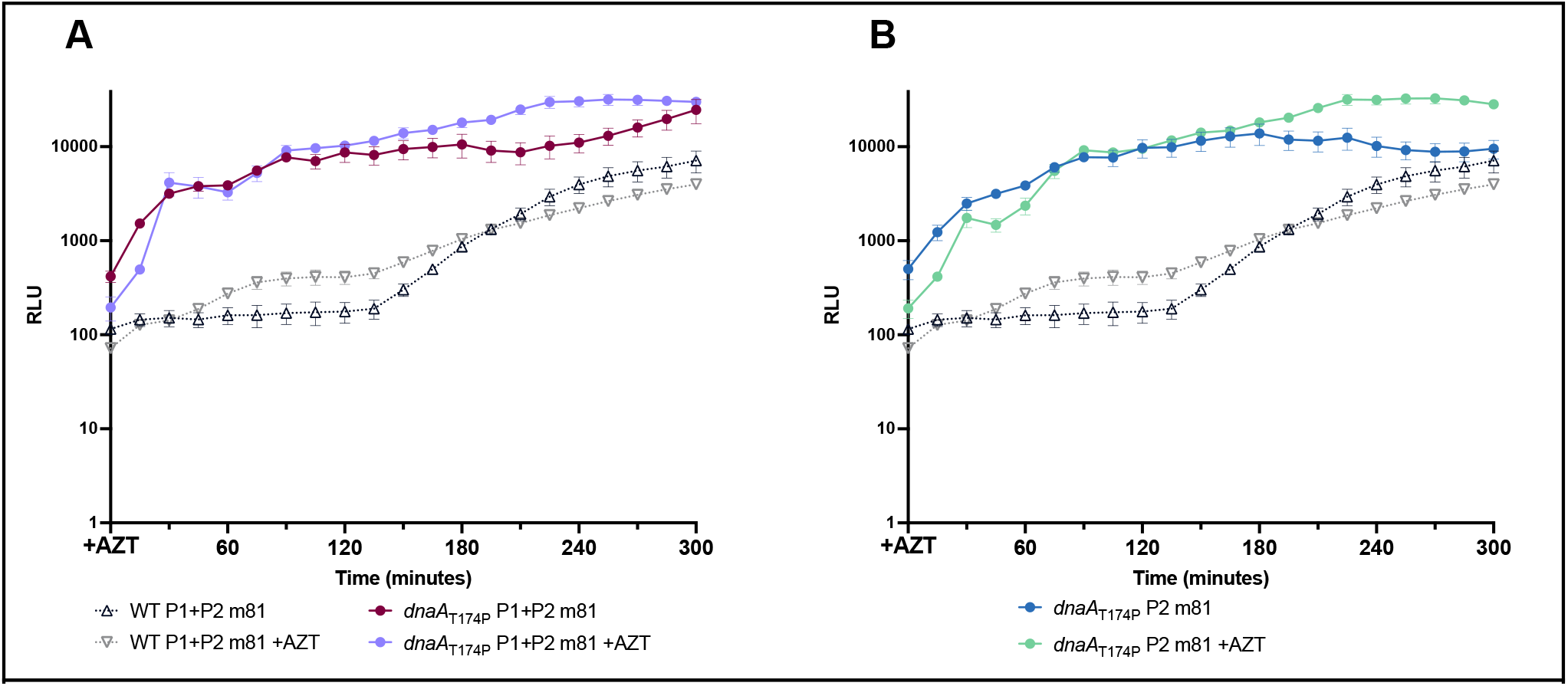
Genetic epistasis of DnaA T174P and DnaA box −81 mutations on iraD promoter expression. A. DnaA Box-81 effects on iraD P1 and P2 B. DnaA Box-81 effects on P2 promoter

Because expression of both P1 and P2 appeared to be positively regulated by the stringent response, we tested expression in strains deficient for *sspA*, “<u>s</u>tringent <u>s</u>tarvation <u>p</u>rotein”, a transcription factor induced by the stringent response [29]. SspA had a strong and complex effect on *iraD* expression, with its strongest effects on P2 (Fig. 7). Consistent with a role in transcriptional activation at the locus, functional SspA promotes higher constitutive expression of *iraD.* However, after sublethal treatment with AZT, functional SspA seems to reduce expression, especially at later time-points. Because of these contrary effects, in *sspA* mutants, the magnitude of DNA-damage induction of *iraD* is much more dramatic in *sspA* than wt strains, for example with 50-60 fold induction vs. 4-6 fold at 150 minutes post-AZT treatment. The genetic effects of SspA are independent of DnaA (Figure 7D), with the SspA protein exerting a positive effect on constitutive expression and a negative effect on AZT-induced expression in a mutant of the DnaA box at −81.

**Figure 7.**
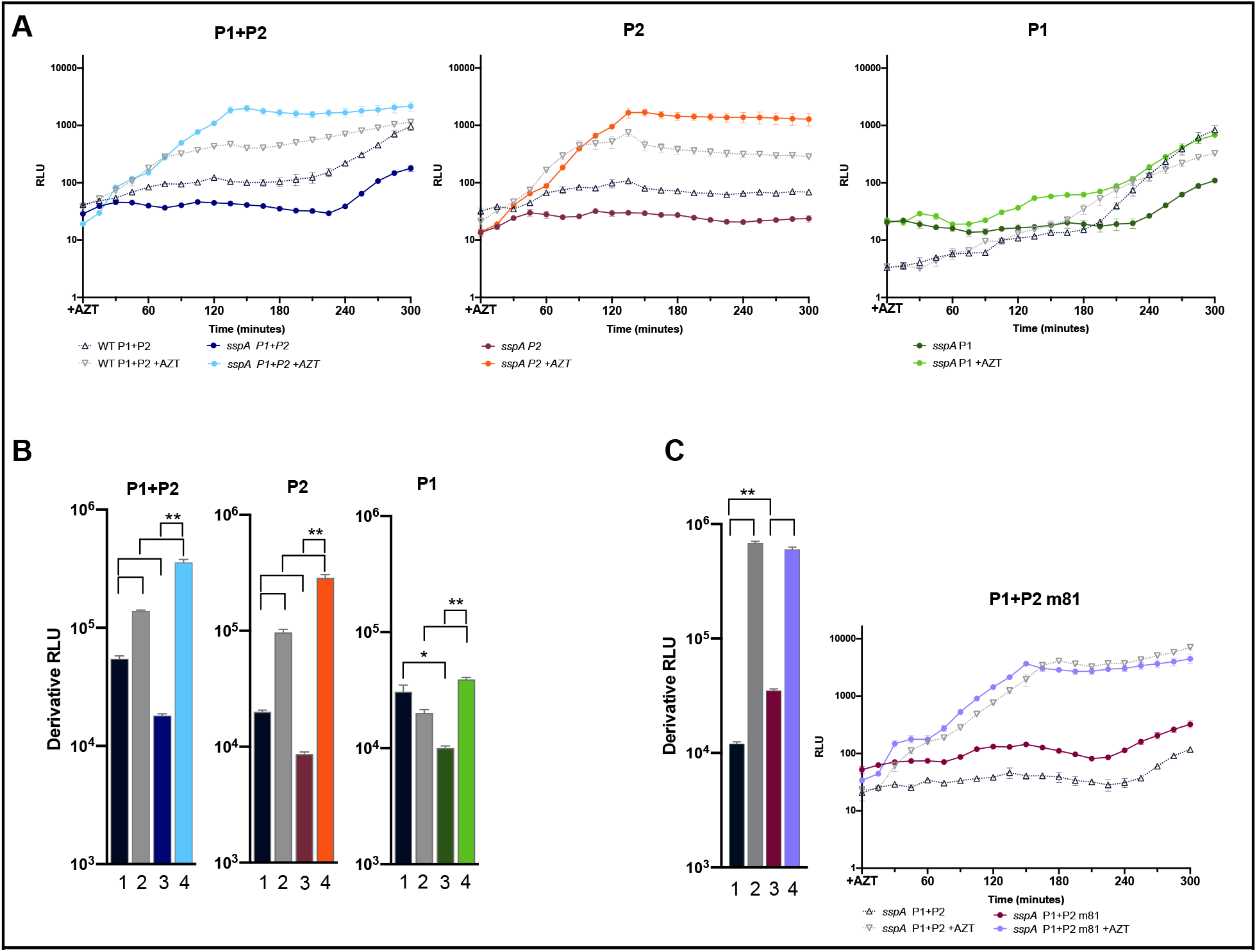
Effects of SspA on IraD expression. A. Expression of luciferase fusions to the 672 bp *iraD* promoter region. RLU is the luminescence normalized to the OD600 as measured during growth of the culture in a plate reader. Note that this is a log scale plot. “P1+P2” carries both intact promoters, “P2” has an inactivated P1 at the −10 region 1; “P1” has inactivated P2 −10 sequence. Open black symbols indicate wt expression; filled colored symbols indicate expression in *sspAΔ* mutants. The darker color indicates constitutive expression, the lighter is that induced by AZT added at 0 min.B. Quantitation of area under the curves in A. C. Effect of DnaA box m-81 in combination with *sspA*. Quantitation at left and RLU expression over time at the right.

## DISCUSSION

### Incomplete replication/DNA damage

The SOS response after DNA damage in *E. coli* up-regulates at least 40 genes, many of which function to sustain viability [11, 43, 44]. The inducing signal for the SOS response is the formation of a RecA filament on persistent single-strand DNA, a structure that promotes the autocleavage of the LexA repressor protein [reviewed in 45, 46]. Formation of the RecA filament also initiates the process of homologous recombination that can fill ssDNA gaps present in DNA. The replication inhibitor AZT, like UV irradiation [47], is a strong inducer of the SOS response and homologous recombination, with the filament loaded by the RecFOR mediator system [48, 49]. Some of the functions induced by the SOS response can be deleterious, including the error-prone, Y-family DNA polymerases, and often the replication fork needs to be re-assembled, leading some to speculate that its full induction may be delayed and called into play only for very persistent ssDNA [50].

It has been known for some time that there are DNA damage-inducible genes that are not regulated by the LexA/RecA system. The molecular details of their regulation and how induction is signaled remains unclear. Less is known about this group, largely because their induction is smaller in magnitude and slower than those induced by the SOS-response. These genes include members that process DNA or nucleotides, including *dnaA* (replication initiator) and its downstream genes, *dnaN* (replication clamp), and *recF* (homologous recombination mediator) as well as, *nrdAB* (nucleotide reductase), *ssb* (single-strand DNA binding protein), *phr* (photolyase), *dnaE* (DNA polymerase III, alpha subunit), *dnaQ* (DNA pol III, epsilon proofreading exonuclease), *dnaB* (replication fork helicase), *recQ* (DNA helicase) and *iraD* (RpoS regulator), the subject of this study [4, 11,51-54].

We show here using promoter fusions that IraD is regulated by DnaA, as are two other SOS-independent genes, *dnaA* (and its operon) and *nrdAB.* Genetic effects are consistent with DnaA-ATP acting as a transcriptional repressor, dependent in part at a DnaA box sequence 81 nt upstream. Because purified DnaA protein binds to the *iraD* promoter region in vitro, we think that this regulation is direct. DnaA appears to regulate both of *iraD’s* promoters, P2, the primary promoter during exponential growth and P1, the primary promoter in late log/stationary phase and the box 81 site is downstream from both promoters. Expression from both P1 and P2 is induced after treatment with sublethal doses of the replication inhibitor AZT; in *dnaA* and *dnaN* mutants that reduce levels of DnaA-ATP, AZT inducibility is largely lost and expression remains at the high constitutive rate.

Through an interaction with its homolog, Hda, and the replication processivity clamp, DnaA is inactivated, blocking its ability to re-initiate replication [55]. Although RIDA is usually considered only in its role in regulating replication initiation, our work and others [14] show that RIDA can also inactivate DnaA’s function as a transcriptional repressor. Among transcription factors, DnaA is unusual in that it self-assembles as a dynamic oligomeric filament on DNA [17, 56, 57], and it is impossible to predict binding sites by sequence alone, although candidate sites can be identified through consensus [15]. A putative DnaA box at −81 is required for repression of both P1 and P2 in vivo. How repression by DnaA may be relieved by DNA damage is not clear, although we raise the possibility that after a block to replication, there may be an accumulation of free DnaN clamps that bind Hda and inactivate DnaA by stimulating its ATPase activity. Whether expression of other DnaA-regulated genes such as *mioC* [58], *guaB* [59], and *polA* [60], are influenced by DNA damage remains to be determined. As an interesting parallel, DnaA also controls an SOS-independent damage response in *Bacillus subtilis* [61]. However DnaA repression of DNA-damage inducible genes is relieved, we suspect that the inducing signal is more immediate after a replication block and possibly more transient in nature than the SOS response.

### Nutrition

The nutritional state of the cell regulates *iraD* expression by several mechanisms. Promoter P1 is induced late log phase, with the accumulation of ppGpp, as part of the stringent response to various starvation conditions [28]. We show here that *iraD* expression is also regulated by stringent starvation protein, SspA, one of the most abundant proteins induced by the stringent response [30]. Constitutive expression of both *iraD* promoters seems to be enhanced by SspA, consistent with the role of the protein as a transcriptional activator. AZT induction of promoter P1, but not P2, seems to be negatively regulated by SspA, since expression is enhanced in *sspA* mutants. Whether any of these effects is direct remains to be determined. DNA damage inducibility of *iraD* is also enhanced by the SOS-noninducible *lexA3* allele; in this case, the simplest interpretation is that the DNA damage signal persists longer, affecting *iraD* indirectly. Since constitutive expression of a construct lacking the putative DnaA box at −81 is still reduced by *sspA*, SspA likely does not act through an effect on DnaA negative regulation. Likewise, although the DNA damage-inducible *nrdAB* promoter is regulated by DnaA quite similarly, it is not as strongly affected by SspA.

The replication cycle in bacteria is regulated at the level of initiation, from a single origin of replication. Because it takes approximately 50 minutes to replicate the *E. coli* chromosome, forks may initiate under favorable nutritional conditions but encounter starvation later, endangering complete replication of the entire chromosome. The last round of replication before stationary phase/quiescence is likely the most problematic. For factors involved in replication fork repair, it therefore makes sense for the cell to integrate a signal of replication gaps with that of a downturn in nutrition to induce their expression. IraD regulation and consequently RpoS is likely such a response.

What factors that sustain tolerance of replication fork stress are regulated by IraD and RpoS remains to be fully determined. The gene for Exonuclease III, the major apurinic/apyrimidinc endonuclease and 3’ to 5’ dsDNA exonuclease of *E. coli* is required for AZT tolerance in vivo [48] and is induced by RpoS [62]. Important for tolerance of oxidative stress, expression of *katE* (catalase/hydroperoxidase HPII) is induced by RpoS [63], as is *dinB*, translesion DNA polymerase IV [64]. Upon entry to stationary phase, expression of *recN* (the gene encoding the processivity clamp) and *recF* (recombination mediator protein) are upregulated by RpoS, independent of the *dnaA* promoter [65].

## MATERIALS AND METHODS

### *E. coli* K-12 strains and growth

All strains used are isogenic with STL18703 (*rph*+ derivative of MG1655 [66]) and were constructed by P1 transduction [67], selecting for a linked antibiotic resistance marker. These include STL22918 (*lexA3 malF*::Tn10), STL23195 (*sspA*ΔFRT *kan*), STL23334 (*dnaA*-T174P miniTn10*cat*), STL23379 (*dnaN*-G157C miniTn10*cat*), and STL23377 (*dnaA*-A345S miniTn10*cat*). Cultures were grown at 37°C in Luria-Bertani (LB) medium supplemented with appropriate antibiotics kanamycin (20 μg/ml), chloramphenicol (15 μg/ml), ampicillin (100 μg/ml), and/or tetracycline (10 μg/ml). LB [67] Lennox formulation was used for standard growth media. Plate media included the addition of Bacto-agar at 2%.

### Plasmid constructions

Plasmid pDEW201-PiraD was constructed from pDEW201 [53] as previously described [28] using primer pair 5’-GGGGACAAG TTTGTACAAA AAAGCAGGCT TCGAAGGAG ATAGAACCAA TAGTTACCTT TTATGAGTAT TTC-3’ and 5’-GGGGACCACT TTGTACAAGA AAGCTGGGTC TTTGCGCACT CCTGACGTTT AGCA A-3’. Plasmids pDEW201-P1iraD and pDEW201-P2iraD were constructed as follows. PCR products were generated by Phusion (New England Biolabs) PCR using pDEW201-PiraD as template and the primer pairs 5’-GGGGCGGCCG TTGCTGACAG GATTCAGGCC TGTCTC-3’ and 5’-GGGGCGGCCG AAACCTTACT TGCCTATGAA TATCTA-3’ for pDEW201-P1iraD and 5’-GGGGGGCCCA ATAATGCCTG TGAATGGTAT TTTTG-3’ and 5’-GGGGGGCCCG ACCTCAATAT AGCAACATCA AATTC-3’ for pDEW201-P2iraD. The resulting PCR products were restriction digested with EagI (New England Biolabs) for pDEW201-P1iraD and ApaI (New England Biolabs) for pDEW201-P2iraD according to the manufacturer’s instructions. The digest products were then ethanol precipitated and self-ligated at room-temperature overnight using T4 DNA Ligase (New England Biolabs). The resulting PCR products were restriction-digested with EagI (New England Biolabs) for pDEW201-P1iraD and ApaI (New England Biolabs) for pDEW201-P2iraD according to the manufacturer’s instructions. The digest products were then ethanol precipitated and self-ligated at room-temperature overnight using T4 DNA ligase. DnaA Box −81 mutants of pDEW201-PiraD, pDEW201-P1irad and pDEW201P2iradD were generating similarly using primers 5’ atcgtcaagc ttgccattcc cttttgttca ttac (AF99) and digestion with HindIII.

### Luciferase expression assays

Luminescence and OD600 were measured using BioTek Cytation 1 Plate Reader and Costar 96 Well Assay Plate (treated polystyrene, black plate, clear bottom). Colonies were inoculated in LB media in tubes, shaking until OD600 = 0.5 after which they were diluted 1:100 in LB and grown again to ensure log phase growth. In the 96 well plates, cells were diluted 1:100 and grown for 2 hours, before being treated with 1.25 ng/mL AZT. Bioluminescence was measured and normalized to the OD600 yielding relative luminescence units (RLU) every 15 minutes and data are averages of 4 independent replicate cultures.

### RpoS Western blots

RpoS steady-state levels during growth were determined by Western blot analysis [4]. Protein was precipitated following 30 minutes of incubation on ice in 20% ice-cold trichloroacetic acid (TCA), and the pellet was resuspended in sodium dodecyl sulfate buffer.

Quantitation was performed by calculating the integrated density using ImageJ (NIH) and normalizing the values to control bands and reporting the resulting value relative to the value obtained for wild-type.

### His6-tagged DnaA purification

His-tagged DnaA was purified from DB3.1 cells harboring the overexpression plasmid pCA24N-dnaA [68] using a two-stage purification process. Stage 1 consisted of affinity purification with modifications to the method of [69] and Stage 2 consisted of a native purification protocol adapted from [15]. Fresh pCA24N-dnaA harboring transformants of DB3.1 were inoculated into 50 mls of 2xYT media (16 g/L tryptone, 10 g/L yeast extract, and 5 g/L NaCl) supplemented with 15 μg/ml chloramphenicol and incubated with constant shaking overnight at 37°C.

The following day this culture was diluted 1:20 in fresh pre-warmed 2 x YT media supplemented with 15 μg/ml chloramphenicol. This culture was incubated with constant shaking aeration at 37°C until the cell density reached approximately 0.8-0.9. Isopropyl-β-D-thiogalactoside (IPTG) was added to the culture to induce protein expression to a final culture concentration of 1 mM. The cultures were then incubated at 37°C with continued shaking for another 2-3 hours.

Cells were harvested from the culture by low-speed centrifugation at 4300 rpm in a swinging bucket rotor. The cell pellet was resuspended in 0.01 vol (relative to original culture volume) of ice-cold Binding Buffer (20 mM sodium phosphate pH 7.8, 500 mM NaCl, 5 mM imidazole), frozen in a dry-ice/ethanol bath for 3-5 minutes and stored at −80°C until further processing.

The frozen cell paste was thawed at room temperature and placed on ice. Lysozyme was added to a final concentration of 0.1 mg/ml, mixed, and incubated on ice for 30 min. The paste was then subjected to three cycles of heat shock at 42°C, followed by rapid cooling in a dry ice/ethanol bath. Finally, MgCl_2_ was added to a final concentration of 1 mM along with addition of DNase and RNase mixtures and the paste was incubated on ice for an additional 15 minutes. The cell lysate was cleared by centrifugation at 18500 rpm in a Sorvell SS34 rotor for 60 minutes at 4°C.

A nickel resin column was equilibrated by application of 5 volumes of water followed by10 volumes of Binding Buffer. The cleared cell lysate supernatant was batch-incubated with the column resin for 15-30 minutes with gentle agitation at 4°C. The lysate was then cleared from the column by gravity-flow. The column was then washed with three rounds of 3 volumes Wash Buffer 1 (20 mM sodium phosphate pH 7.8, 500 mM NaCl, and 40 mM imidazole), 8 volumes Urea Wash Buffer (20 mM sodium phosphate pH 7.8, 500 mM NaCl, 5 mM imidazole, and 7 M urea), and finally three rounds of 3 volumes Wash Buffer 2 (50 mM Hepes-KOH pH 7.6, 100 mM potassium glutamate, 10 mM magnesium acetate, and 20% sucrose). Protein was eluted from the column by application of five 1 mL treatments of Elution Buffer (45 mM Hepes-KOH pH 7.6, 750 mM potassium glutamate, 10 mM magnesium acetate, 500 mM imidazole, and 20% sucrose). Protein containing fractions were pooled, and dialyzed against Dialysis Buffer (45 mM Hepes-KOH pH 7.6, 750 mM potassium glutamate, 10 mM magnesium acetate, 1 mM DTT, 0.5 mM EDTA, and 20% sucrose) for 1 hour with stirring agitation for 1 hour at 4°C. Aggregated protein was cleared centrifugation for 30-60 minutes at 18500 rpm at 4°C and aliquots were immediately frozen and stored at −80°C until Stage 2 processing.

For Stage 2 purification, frozen protein aliquots were thawed completely on ice, pooled, and brought to a final volume of 2.5 mls with LG Buffer (45 mM Hepes-KOH pH 7.6, 100 mM potassium glutamate, 10 mM magnesium acetate, 1 mM DTT, 0.5 mM EDTA, and 20% sucrose). This solution was applied to a PD-10 desalting column previously equilibrated with 25 mls LG Buffer and then eluted by application of 3.5 mls LG Buffer to the column. The eluted fraction was then applied to a HiTrap HP SP ion exchange column that was previously equilibrated with LG Buffer. The column was washed with 5-10 volumes of LG Buffer and the protein was eluted with a step-wise 0.1-1 M potassium glutamate gradient prepared in LG Buffer. Fractions were collected and aliquots frozen and stored at −80°C.

### Electrophoretic mobility shift assays (EMSA)

EMSA assays were performed as previously described [15] with the following modifications. The DNA used for each reaction was generated by PCR and consisted of the 672 basepair iraD intergenic region. His-tagged DnaA was incubated with 10 nM of DNA in binding buffer (20 mM Hepes pH 8.0, 1 mM EDTA, 8 mM DTT, 5 mM MgOAc, 10% glycerol, 0.1 mg/ml BSA, and 1 mM ATP) at 16°C for 30 minutes. Electrophoresis of the reactions was performed in 1.5% agarose in 0.5X TBE at 4°C. The gel was stained in 1X Sybr Green I in electrophoresis buffer and the DNA visualized using UV according to the manufacturer’s instructions. Densitometry was performed using ImageJ software (NIH). Least-square fitting of the binding isotherm and determination of binding constants are statistics were performed using Graphpad Prism software.

## Acknowledgments

We thank Jon Beckwith for providing *dnaA* and *dnaN* mutant strains. This work was supported by NIH grants R01 GM51753 to STL and T32 GM001722 to THS.

